# Respiration and gas exchange under negative pressure breathing during simulated microgravity

**DOI:** 10.1101/2024.04.08.588548

**Authors:** Yury S. Semenov, Aleksandra A. Gorbunova, Julia A. Popova, Alexander I. Dyachenko

## Abstract

The research considers the effect of inspiratory negative pressure breathing (NPBin, -20 cmH_2_O relative to barometric pressure) on respiration and gas exchange in healthy humans under various conditions: in the sitting and supine positions as well as during simulating the physiological effects of a long stay in conditions of microgravity (dry immersion and head-down bed rest). Under NPBin, respiratory rate significantly decreased (by 7 min^-1^ on average, in some volunteers up to two or three cycles per minute), tidal volume increased (by an average of 0.3 l), while minute ventilation decreased (on average by 1.8 l/min), and respiratory exchange ratio increased (on average by 0.09). The response of respiratory and gas exchange parameters to NPBin in all the considered conditions was almost the same (there were no significant differences in parameters changes induced by NPBin between the conditions). Despite the decrease in minute ventilation, the results indicate CO_2_ washout from the body under NPBin.

## 1. Introduction

Negative pressure breathing is a mode of breathing when pressure below atmospheric is created and maintained in the airways. Pressure reduction can be implemented either throughout the entire respiratory cycle (NPB) or in its certain phases. In this paper we consider additional inspiratory negative pressure breathing (NPBin), the additional rarefaction in the airways is created during the inspiratory phase only. During the remaining phases of the respiratory cycle airway pressure does not change relative to normal breathing.

In the early 1990s Tikhonov et al. have proposed NPB as a method to prevent undesirable physiological consequences of being in microgravity during spaceflight (Tikhonov et al., 1991). According to experimental data (Baranov et al., 2000), NPB -20 cmH_2_O (hereinafter all pressure values are given relative to barometric pressure) allows to decrease intrathoracic pressure by about -10 cmH_2_O. This value is approximately equal to the decrease in pressure in blood vessels at the level of chest when moving from a horizontal to a vertical position. Thus, the use of NPB makes it possible to partially simulate physiological effects of verticalization.

NPBin is also considered as a technique to prevent headward fluid shift caused by microgravity (Popova et al., 2022). Moreover, NPBin is used in terrestrial medicine to increase the effectiveness of resuscitation measures as well as to maintain the proper values of central and cerebral hemodynamic parameters in case of blood loss and other hypotonic conditions (Convertino, 2019; Lurie et al., 2001, 2004; Rickards, 2019). However, changes in the pattern of breathing and gas exchange caused by NPBin remain insufficiently studied.

**The aim of our research** was to study changes in respiratory pattern and gas exchange caused by NPBin in healthy subjects at rest and under conditions of dry immersion (DI) (Navasiolava et al., 2011; Tomilovskaya et al., 2019) and head-down bed rest (HDBR) (Hargens and Vico, 2016; Watenpaugh, 2016), which are on-ground models for simulating physiological effects of prolonged exposure to microgravity. We additionally examined changes in the response to NPBin with changing in body position (supine or sitting), since the body position has a profound effect on the distribution of blood flow and ventilation in the lungs (Prisk, 2000), and, apparently, may affect the result of using NPBin.

A number of studies that considered NPB also took into account body position (Baranov et al., 2001, 2003; Tikhonov et al., 2003). However, in these studies a change in body position was considered primarily as a way to cause redistribution of blood in the cranial direction, which to be minimized with NPB. The response of breathing pattern and gas exchange to NPB or NPBin had not been studied in detail.

## 2. Materials and methods

### 2.1 Design

Two different groups of volunteers participated in our study: one group in DI and another in HDBR experiment, respectively. All subjects underwent a medical history and physical examination by a physician to ensure that they had no previous or current medical conditions that might preclude their participation. In accordance with the Declaration of Helsinki, all subjects gave their written informed consent to participate in the study. The study was approved by the Bioethics Commission of the Institute of Biomedical Problems of the Russian Academy of Sciences (protocols № 299 of 30.05.2012 and № 302 of 25.07.2012).

In each experiment, the analysis of NPBin effect on parameters of external respiration and gas exchange was carried out using the same measuring equipment as well as the same measurement protocol, the only difference was the experimental condition (DI, HDBR, or body position). Therefore, we used a unified approach to processing the data obtained in the experiments and considered the results together.

NPBin was implemented using a face mask with a valve box, which separates the inspiratory and expiratory flows. L-port three-way valve and a spring-loaded valve were additionally installed in the inspiratory circuit. The spring-loaded valve (NPBin-valve) opens when a certain pressure difference between the inspiratory line and an atmosphere is reached. This valve is structurally similar to a standard gas reducer. The adjustable spring preload of the valve sets the pressure in the inspiratory line. The three-way valve was used to disconnect spring-loaded valve from the system during free breathing. Since the inspiratory and expiratory flows were separated, additional dead volume was created by the facemask only and remained unchanged when switching from free breathing to NPBin and vice versa. NPBin-valve was set to -20 cmH_2_O.

Volunteers were acquainted with the measuring equipment and NPBin several days before the start of the study. Each of the study series was divided into three stages (there were no pauses between them): free breathing (but through the respiratory equipment) for 15 minutes, the NPBin stage or continuation of free breathing in the control series (for convenience also called “NPBin” in the text and table below) for 25 minutes, free breathing after NPBin (also through breathing equipment) for 15 minutes.

In the first experiment, a 15-hour HDBR with an angle of -15° to the horizon was used to simulate the physiological effects of a long stay in microgravity. Measurements were performed at the end of the 14th or at the beginning of the 15th hour of HDBR. Eight healthy men of average build aged 22 to 35 years participated in the experiment. The experiment included two series (the HDBR-NPBin series and the HDBR-CONTROL series) with identical procedures schedule; the only difference is there was no actual NPBin in HDBR-CONTROL series. Each of the eight volunteers participated in both series. The sequence of series for an individual volunteer was chosen randomly, 4 volunteers first had the HDBR-NPBin series, other 4 volunteers – HDBR-CONTROL. The duration of the rest between series for an individual volunteer was several days or more.

The second experiment (3-day DI) involved 13 volunteers (healthy men of average build aged 19 to 28 years). Each of the 13 volunteers participated in six series including a control one (Immersion-BDC SUPINE, Immersion-BDC SITTING, Immersion-DI, Immersion-RECOVERY SUPINE, Immersion-RECOVERY SITTING, Immersion-CONTROL).

The Immersion-BDC SUPINE and Immersion-BDC SITTING series were performed on the same day, a few days before DI, with a rest period between series of at least two hours. Series differed only in the position of the volunteer. The sequence of body position for an individual volunteer was chosen randomly.

The Immersion-DI series was carried out on the second day of DI. During the measurements, the volunteer was in an immersion bath.

The series Immersion-RECOVERY SUPINE and Immersion-RECOVERY SITTING were carried out on the second day after the end of DI, the rest period between series was at least two hours, this pair of series repeated Immersion-BDC SUPINE and Immersion-BDC SITTING series.

In the pairs of series Immersion-BDC SITTING/SUPINE and Immersion-RECOVERY SITTING/SUPINE, half of the volunteers first participated in the SUPINE series, then SITTING, and the other half, on the contrary, first participated in the SITTING series. For each volunteer the sequence of positions in Immersion-RECOVERY series was opposite to the sequence from the Immersion-BDC series.

The Immersion-CONTROL series was carried out a month after the end of DI. The volunteer was lying horizontally on his back during measurements. This series completely repeated the series corresponding to the supine position, except the NPBin-valve was not activated.

### 2.2 Measurements

Continuous measurements of inspiratory flow, pressure under the mask (hereinafter we denote it as mouth pressure, Pm), and expiratory concentration of O_2_ and CO_2_ were performed throughout each series (MPR6-03 Monitor, Triton Electronic Systems Ltd., Russia). Both “minute” parameters of ventilation and gas exchange (O_2_ uptake 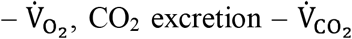, respiratory exchange ratio – R, respiratory minute volume – MV) and breath-by-breath parameters (tidal volume – V_T_, end-tidal partial pressures of O_2_ and 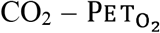 and 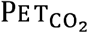) were calculated. Moreover, breathing pattern characteristics were obtained: duration of respiratory cycle (T, f_R_=1/T – respiratory rate), duration of inspiration (T_I_), duration of expiration (T_E_), duration of the pause between inspiration and subsequent expiration (Tp_1_), duration of the pause between expiration and subsequent inspiration (Tp_2_). Also we calculated the work associated with inspiration (W) as an integral of the product of inspiratory flow and Pm, the integration boundaries coincide with the edges of inspiration. Gas exchange and ventilation parameters were converted to standard reference conditions (BTPS and STPD). Peak rarefaction during inspiration and peak pressure during expiration, as well as the average value of Pm throughout the stage, were calculated from the recordings of Pm (the average pressure value was calculated without distinguishing inspirations and expirations).

To estimate gas exchange in tissues a transcutaneous O_2_ and CO_2_ tension monitor 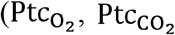; TCM4, Radiometer, Denmark) was used. The sensor was located on skin surface in the left subclavian region. Also pulse oximetry (TCM4, Radiometer, Denmark) was used to continuously measure arterial blood oxygen saturation (SpO_2_).

In addition, O_2_ and CO_2_ tension in arterialized capillary blood were measured (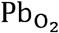, 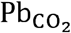). Four blood samples were taken: before the start of “NPBin” stage, at the 12th and 25th minutes of NPBin, and at 15th minute after the end of NPBin. Samples were analyzed with ABL80 (Radiometer, Denmark). During data processing, the results of the analysis of two samples taken at “NPBin” stage were averaged as there was no statistically significant difference between them (Wilcoxon signed-rank test, p>0.05).

### 2.3 Data analysis

The measured data for each of the series were processed according to a common template. For each recording (i.e. for an individual subject), the average value of the parameter per stage (“before NPBin”, “NPBin”, “after NPBin”) was calculated. Also we calculated the difference between the average values of the parameter for current stage and stage “before NPBin” for each subject. Group descriptive statistics were obtained for average-per-stage values and for the differences. Wilcoxon signed rank test was used for pairwise comparisons of the stages within the series, or the same stages in different series. Comparison of stages within the series was used to reveal NPBin effects. Comparison of the same stages in different series was used to find changes in NPBin-induced reactions caused by body position or DI, this comparison was applied only to the differences.

## 3. Results

### 3.1 Breathing pattern and expired gas analysis

Switching from free breathing to NPBin or backwards, respiratory parameters were changing over a period from several respiratory cycles to several minutes (the duration of the transition varies greatly in different volunteers), but then the parameters remained stable until NPBin applied. The results presented below were obtained for sections of recordings that did not include two-minute segments around the moment of turning on or turning off the NPBin mode.

All volunteers demonstrated normal values of respiratory and gas exchange parameters both before and after the use of NPBin.

Volunteers demonstrated the same type of response to NPBin in any of the series, regardless of the conditions (supine or sitting position, HDBR, DI). Respiratory rate (f_R_) decreased and V_T_ increased; MV decreased or did not change. However, the magnitude of the change in f_R_ and V_T_ varied greatly among different volunteers, in one person f_R_ and V_T_ could remain virtually unchanged, while in another f_R_ decreased to 2-3 cycles per minute, and V_T_ increased almost to its maximum possible value. The response to NPBin was reproducible in each of the volunteers in different series of the study. After the end of NPBin, parameters returned to their original values in several minutes or less (in most volunteers it took only several respiratory cycles).

The maximum Pm value at any of the stages in any of the study series did not exceed 1.5 cmH_2_O. The maximum rarefaction under the mask at the stages of free breathing in any of the study series did not exceed 1.0 cmH_2_O. The rarefaction during inspiration under NPBin in any of the study series was in the range of 18-22 (median 19.2 cmH_2_O). The average Pm over the entire “NPBin” stage ranged from -2.9 to -6.4 (median -4.7 cmH_2_O), sign “-” indicates rarefaction. The average Pm over the entire stage at the stages of free breathing was zero. A large dispersion of average-over-stage Pm values was caused by a large individual variation of NPBin-induced changes of breathing pattern, mainly, by the duration of the pause between expiration and subsequent inspiration (Tp_2_). The work associated with inspiration (W) under NPBin was also characterized by a large individual scatter, W ranged from 10 to 44 (median 14 l·cmH_2_O). During free breathing W was 0.3 l·cmH_2_O.

Table 1 shows data for a group of volunteers, the results are presented in two versions: the value of the parameter and its change over the series relative to the individual baseline value (average-per-stage value observed in a volunteer at “before NPBin” stage).

**Table 1.** Respiratory and gas exchange parameters at stages of the series and their changes induced by NPBin.

|  | Average-per-stage values of parameters |  |  | Differences in parameters between the stages |  |
| --- | --- | --- | --- | --- | --- |
|  | before NPBin | NPBin | after NPBin | NPBin – before NPBin | after NPBin – before NPBin |
| Series: HDBR-NPBin |  |  |  |  |  |
| PET<sub>CO<sub>2</sub></sub> (mmHg, BTPS) | 29.3 [28.2; 30.0] | 34.0 [30.6; 35.0] | 26.8 [25.6; 28.5] | 4.6* [3.2; 5.3] | -1.1 [-3.6; 0.9] |
| PET<sub>O<sub>2</sub></sub> (mmHg, BTPS) | 112.5 [110.6; 114.3] | 107.7 [106.4; 109.2] | 110.6 [109.3; 113.7] | -4.8* [-5.7; -3.9] | -1.8 [-4.1; 1.6] |
| R | 0.775 [0.761; 0.882] | 0.892 [0.806; 0.937] | 0.732 [0.692; 0.928] | 0.091** [0.028; 0.129] | -0.064 [-0.072; 0.062] |
| $\dot{V}_{CO_2}$ (l/min, STPD) | 0.20 [0.13; 0.22] | 0.20 [0.17; 0.23] | 0.19 [0.16; 0.21] | 0.02 [0.00; 0.04] | -0.01 [-0.02; 0.02] |
| $\dot{V}_{O_2}$ (l/min, STPD) | 0.25 [0.16; 0.28] | 0.23 [0.19; 0.25] | 0.26 [0.21; 0.28] | 0.00 [-0.03; 0.04] | 0.01 [-0.02; 0.04] |
| MV (l/min, BTPS) | 8.6 [5.9; 9.9] | 7.3 [6.8; 7.9] | 8.5 [7.7; 9.3] | -1.8 [-2.5; 0.4] | -0.5 [-1.4; 0.9] |
| VT (l, BTPS) | 0.49 [0.35; 0.55] | 0.85 [0.68; 2.28] | 0.52 [0.43; 0.66] | 0.44** [0.14; 1.85] | 0.1 [-0.03; 0.18] |
| fr (min<sup>-1</sup>) | 17.0 [14.9; 20.1] | 8.4 [2.4; 13.1] | 16.5 [12.3; 19.6] | -9.5** [-12.2; -7.6] | -1.7 [-3.2; 0.6] |
| Ti (s) | 1.3 [1.2; 1.4] | 2.1 [1.6; 2.1] | 1.4 [1.2; 1.8] | 0.7* [0.2; 0.8] | 0.1 [-0.1; 0.4] |
| Te (s) | 1.6 [1.6; 1.8] | 2.4 [2.1; 4.2] | 1.7 [1.4; 2.3] | 0.9* [0.7; 2.4] | 0.2 [-0.1; 0.4] |
| Tp<sub>2</sub> (s) | 0.2 [0.1; 0.6] | 1.4 [0.6; 15.3] | 0.8 [0.1; 1.5] | 1.6* [0.6; 14.5] | 0.0 [0.0; 1.2] |
| Ptc<sub>CO<sub>2</sub></sub> (mmHg) | 41.0 [40.3; 43.8] | 39.3 [38.1; 41.0] | 39.0 [38.6; 40.5] | -2.6* [-3.4; -1.6] | -1.7* [-2.9; -1.3] |
| Pb<sub>CO<sub>2</sub></sub> (mmHg) | 39.0 [38.0; 40.0] | 41.3 [40.0; 42.5] | 40.0 [39.5; 44.0] | 1.8* [1.0; 3.0] | 0.5 [-0.5; 3.0] |
| Series: HDBR-CONTROL |  |  |  |  |  |
| PET<sub>CO<sub>2</sub></sub> (mmHg, BTPS) | 30.0 [27.7; 31.7] | 28.3 [26.9; 30.4] | 28.7 [25.9; 29.5] | -0.8* [-2.2; -0.3] | -1.7* [-2.4; -1.2] |
| PET<sub>O<sub>2</sub></sub> (mmHg, BTPS) | 111.2 [106.9; 113.8] | 112.4 [109.7; 114.0] | 112.5 [109.2; 115.9] | 1.1* [0.6; 1.8] | 2.0* [0.8; 3.4] |
| R | 0.774 [0.767; 0.854] | 0.796 [0.747; 0.876] | 0.783 [0.746; 0.881] | 0.000 [-0.020; 0.022] | 0.003 [-0.017; 0.009] |
| $\dot{V}_{CO_2}$ (l/min, STPD) | 0.19 [0.15; 0.22] | 0.18 [0.14; 0.21] | 0.17 [0.13; 0.21] | -0.02 [-0.02; 0.00] | -0.01 [-0.02; -0.01] |
| $\dot{V}_{O_2}$ (l/min, STPD) | 0.23 [0.16; 0.29] | 0.21 [0.15; 0.27] | 0.20 [0.15; 0.27] | -0.02* [-0.02; 0.00] | -0.02 [-0.03; -0.01] |
| MV (l/min, BTPS) | 8.8 [6.3; 9.6] | 8.2 [6.2; 9.1] | 7.9 [6.0; 9.0] | -0.1 [-0.7; 0.3] | -0.5 [-0.7; 0.1] |
| VT (l, BTPS) | 0.50 [0.43; 0.56] | 0.45 [0.34; 0.56] | 0.46 [0.39; 0.55] | 0.00 [-0.13; 0.00] | -0.03 [-0.06; 0.00] |
| fr (min<sup>-1</sup>) | 16.9 [12.3; 20.6] | 16.3 [14.5; 20.6] | 16.9 [14.8; 20.8] | 0.6 [-0.7; 1.9] | 1.5 [-0.9; 2.4] |
| Ti (s) | 1.3 [1.2; 1.6] | 1.3 [1.1; 1.5] | 1.4 [1.1; 1.5] | 0.0 [-0.2; 0.0] | -0.1 [-0.2; 0.0] |
| Te (s) | 1.8 [1.5; 2.1] | 1.7 [1.4; 2.1] | 1.7 [1.3; 2.1] | 0.0 [-0.3; 0.0] | -0.1 [-0.5; 0.1] |
| Tp<sub>2</sub> (s) | 0.1 [0.1; 0.5] | 0.2 [0.1; 0.7] | 0.1 [0.1; 0.7] | 0.0 [-0.2; 0.0] | 0.0 [0.0; 0.0] |
| Ptc<sub>CO<sub>2</sub></sub> (mmHg) | 40.9 [37.0; 43.2] | 39.9 [37.2; 42.2] | 40.4 [38.1; 41.9] | -0.9* [-1.0; -0.6] | -1.5 [-2.2; 0.0] |
| Pb<sub>CO<sub>2</sub></sub> (mmHg) | 40.5 [39.5; 42.0] | 42.5 [41.0; 43.5] | 42.0 [40.5; 44.0] | 2.8 [-0.8; 4.0] | 1.5 [-1.5; 4.5] |
| Series: Immersion-BDC SITTING |  |  |  |  |  |
| PET<sub>CO<sub>2</sub></sub> (mmHg, BTPS) | 30.0 [28.6; 31.3] | 30.3 [28.1; 33.2] | 29.3 [26.6; 30.9] | 1.5 [-0.9; 3.5] | -0.7 [-1.9; 1.0] |
| PET<sub>O<sub>2</sub></sub> (mmHg, BTPS) | 110.6 [108.8; 113.6] | 111.6 [106.2; 115.0] | 110.8 [105.2; 113.0] | -1.2 [-3.3; 2.6] | -0.8 [-3.9; 0.6] |
| R | 0.814 [0.789; 0.857] | 0.929 [0.849; 0.947] | 0.754 [0.702; 0.801] | 0.078*** [0.059; 0.117] | -0.060* [-0.124; 0.027] |
| $\dot{V}_{CO_2}$ (l/min, STPD) | 0.24 [0.19; 0.27] | 0.24 [0.20; 0.27] | 0.22 [0.17; 0.27] | 0.02 [-0.03; 0.04] | 0.00 [-0.02; 0.02] |
| $\dot{V}_{O_2}$ (l/min, STPD) | 0.26 [0.23; 0.32] | 0.25 [0.23; 0.29] | 0.30 [0.22; 0.35] | -0.02 [-0.05; 0.03] | 0.02 [-0.01; 0.05] |
| MV (l/min, BTPS) | 10.5 [8.4; 11.7] | 9.0 [7.0; 11.4] | 9.6 [7.3; 11.0] | -1.1 [-2.8; 0.9] | -0.5 [-1.3; 0.5] |
| VT (l, BTPS) | 0.66 [0.55; 0.73] | 1.04 [0.75; 1.35] | 0.60 [0.41; 0.83] | 0.31** [0.02; 0.64] | -0.02 [-0.13; 0.08] |
| fr (min<sup>-1</sup>) | 16.7 [13.1; 19.2] | 11.0 [6.7; 13.4] | 17.6 [12.8; 18.3] | -5.9*** [-6.5; -5.2] | -1.1 [-2.2; 1.1] |
| Ti (s) | 1.3 [1.2; 1.5] | 1.5 [1.1; 1.7] | 1.2 [1.1; 1.5] | 0.1 [-0.2; 0.3] | -0.1 [-0.1; 0.0] |
| Te (s) | 1.8 [1.6; 2.3] | 2.3 [1.9; 2.7] | 1.9 [1.5; 2.1] | 0.4* [0.0; 0.7] | 0.0 [-0.2; 0.6] |
| Tp<sub>2</sub> (s) | 0.2 [0.1; 0.9] | 0.2 [0.1; 1.7] | 0.7 [0.1; 1.2] | 0.1 [-0.1; 1.6] | 0.1 [-0.2; 0.9] |
| Ptc<sub>CO<sub>2</sub></sub> (mmHg) | 40.2 [38.2; 43.3] | 36.5 [34.3; 40.0] | 38.6 [35.9; 40.2] | -3.3*** [-7.4; -0.8] | -1.9*** [-4.1; -0.6] |
| Pb<sub>CO<sub>2</sub></sub> (mmHg) | 39.0 [38.0; 40.0] | 38.3 [35.0; 40.3] | 42.0 [40.3; 43.0] | -1.5 [-4.5; 2.3] | 3.0*** [2.0; 3.0] |
| Series: Immersion-BDC SUPINE |  |  |  |  |  |
| PET<sub>CO<sub>2</sub></sub> (mmHg, BTPS) | 31.8 [29.5; 35.6] | 34.3 [31.7; 36.7] | 31.4 [29.0; 33.3] | 2.4 [-0.1; 3.9] | -1.8 [-2.9; 0.9] |
| PET<sub>O<sub>2</sub></sub> (mmHg, BTPS) | 110.9 [108.2; 112.9] | 109.2 [106.1; 111.8] | 107.0 [105.5; 109.8] | -1.2 [-3.6; 1.2] | -2.8 [-5.2; -1.2] |
| R | 0.886 [0.791; 0.977] | 0.995 [0.875; 1.029] | 0.729 [0.707; 0.846] | 0.091*** [0.044; 0.125] | -0.045** [-0.224; -0.007] |
| $\dot{V}_{CO_2}$ (l/min, STPD) | 0.25 [0.20; 0.28] | 0.25 [0.18; 0.27] | 0.20 [0.16; 0.22] | -0.02 [-0.05; 0.03] | -0.05*** [-0.07; -0.01] |
| $\dot{V}_{O_2}$ (l/min, STPD) | 0.27 [0.23; 0.30] | 0.24 [0.22; 0.25] | 0.27 [0.22; 0.29] | -0.04* [-0.08; 0.01] | -0.01 [-0.03; 0.01] |
| MV (l/min, BTPS) | 9.7 [7.8; 11.9] | 8.3 [6.4; 9.6] | 8.5 [7.2; 9.7] | -2.6** [-3.7; -0.8] | -1.3*** [-2.1; -0.7] |
| VT (l, BTPS) | 0.64 [0.53; 0.79] | 0.95 [0.66; 1.46] | 0.55 [0.45; 0.73] | 0.38*** [0.04; 0.79] | -0.06** [-0.10; -0.01] |
| fr (min<sup>-1</sup>) | 17.3 [15.0; 18.3] | 9.3 [5.7; 12.6] | 15.2 [13.4; 18.8] | -7.7*** [-11.1; -4.8] | -1.5* [-2.0; 0.3] |
| Ti (s) | 1.4 [1.2; 1.7] | 1.5 [1.4; 2.0] | 1.5 [1.2; 1.7] | 0.1 [-0.2; 0.6] | 0.1 [-0.1; 0.2] |
| Te (s) | 1.8 [1.6; 2.0] | 2.3 [1.8; 3.9] | 1.9 [1.6; 2.4] | 0.6* [0.3; 1.7] | 0.1 [-0.2; 0.3] |
| Tp<sub>2</sub> (s) | 0.1 [0.1; 0.3] | 0.2 [0.1; 1.7] | 0.2 [0.1; 0.4] | 0.2*** [0.0; 1.5] | 0.0 [0.0; 0.4] |
| Ptc<sub>CO<sub>2</sub></sub> (mmHg) | 40.6 [40.1; 42.5] | 39.1 [36.7; 42.2] | 40.2 [37.2; 42.3] | -2.1*** [-2.9; -1.0] | -1.0 [-3.8; 0.2] |
| Pb<sub>CO<sub>2</sub></sub> (mmHg) | 41.0 [39.0; 42.0] | 39.5 [38.8; 42.5] | 43.0 [42.0; 44.0] | 0.5 [-2.3; 1.5] | 2.0* [0.0; 3.0] |
|  | Series: Immersion-DI |  |  |  |  |
| PET<sub>CO<sub>2</sub></sub> (mmHg, BTPS) | 33.1 [30.3; 35.1] | 35.7 [33.9; 37.4] | 32.1 [29.1; 34.6] | 2.9*** [1.7; 5.1] | -0.2 [-0.7; 1.1] |
| PET<sub>O<sub>2</sub></sub> (mmHg, BTPS) | 108.7 [106.8; 110.6] | 105.7 [104.6; 109.3] | 110.1 [106.3; 112.0] | -2.1 [-5.0; -0.5] | 0.6 [-0.8; 2.2] |
| R | 0.872 [0.782; 0.908] | 0.934 [0.890; 1.039] | 0.857 [0.788; 0.901] | 0.073*** [0.035; 0.137] | -0.010 [-0.030; 0.033] |
| $\dot{V}_{CO_2}$ (l/min, STPD) | 0.23 [0.22; 0.26] | 0.24 [0.22; 0.25] | 0.24 [0.22; 0.25] | 0.00 [-0.02; 0.04] | 0.00 [-0.02; 0.02] |
| $\dot{V}_{O_2}$ (l/min, STPD) | 0.28 [0.26; 0.31] | 0.25 [0.22; 0.27] | 0.28 [0.26; 0.29] | -0.02 [-0.04; 0.00] | -0.01 [-0.02; 0.02] |
| MV (l/min, BTPS) | 9.5 [8.5; 9.9] | 8.1 [6.8; 8.5] | 9.5 [8.7; 10.0] | -1.6*** [-2.6; -0.5] | 0.1 [-0.6; 1.0] |
| VT (l, BTPS) | 0.58 [0.50; 0.64] | 0.66 [0.60; 1.24] | 0.54 [0.50; 0.61] | 0.19* [0.07; 0.71] | -0.01 [-0.10; 0.04] |
| fr (min<sup>-1</sup>) | 16.9 [14.5; 19.1] | 12.5 [5.5; 14.4] | 17.1 [14.6; 20.6] | -6.9*** [-8.7; -3.8] | 0.4 [-0.9; 2.5] |
| Ti (s) | 1.4 [1.3; 1.6] | 1.1 [1.0; 1.6] | 1.3 [1.1; 1.5] | -0.2 [-0.4; 0.3] | -0.1* [-0.2; 0.0] |
| TE (s) | 1.8 [1.6; 2.3] | 1.8 [1.6; 2.6] | 1.9 [1.6; 2.1] | -0.1 [-0.2; 0.5] | -0.2 [-0.3; 0.1] |
| Tp<sub>2</sub> (s) | 0.1 [0.0; 0.3] | 0.6 [0.1; 4.7] | 0.2 [0.1; 0.3] | 0.6* [0.0; 4.5] | 0.1 [-0.1; 0.2] |
| Ptc<sub>CO<sub>2</sub></sub> (mmHg) | 44.4 [41.7; 46.4] | 42.2 [39.1; 46.0] | 41.8 [39.9; 46.2] | -1.0 [-3.0; 0.3] | -1.8 [-3.3; -0.1] |
| Pb<sub>CO<sub>2</sub></sub> (mmHg) | 42.0 [38.3; 43.0] | 41.5 [39.3; 44.3] | 44.0 [42.3; 47.0] | 2.0 [1.1; 2.8] | 4.0*** [3.3; 6.0] |
|  | Series: Immersion-RECOVERY SUPINE |  |  |  |  |
| PET<sub>CO<sub>2</sub></sub> (mmHg, BTPS) | 32.2 [30.1; 33.3] | 34.6 [33.0; 37.5] | 32.9 [30.3; 34.3] | 3.1* [1.1; 4.6] | 0.2 [-0.9; 1.4] |
| PET<sub>O<sub>2</sub></sub> (mmHg, BTPS) | 110.9 [105.6; 112.4] | 107.6 [104.4; 110.7] | 106.6 [103.2; 112.3] | -2.4 [-5.1; 0.2] | -1.8** [-2.6; -0.7] |
| R | 0.867 [0.799; 0.899] | 0.907 [0.885; 1.007] | 0.793 [0.762; 0.895] | 0.093*** [0.069; 0.129] | -0.006 [-0.092; 0.018] |
| $\dot{V}_{CO_2}$ (l/min, STPD) | 0.25 [0.22; 0.26] | 0.24 [0.21; 0.27] | 0.22 [0.21; 0.27] | -0.01 [-0.04; 0.01] | -0.01 [-0.03; 0.00] |
| $\dot{V}_{O_2}$ (l/min, STPD) | 0.28 [0.26; 0.34] | 0.25 [0.22; 0.31] | 0.28 [0.26; 0.34] | -0.04* [-0.06; -0.01] | -0.01 [-0.01; 0.02] |
| MV (l/min, BTPS) | 10.7 [9.7; 11.5] | 7.7 [6.6; 9.6] | 10.0 [8.0; 10.7] | -2.4*** [-4.1; -1.2] | -1.0* [-1.8; -0.1] |
| VT (l, BTPS) | 0.58 [0.53; 0.66] | 0.73 [0.55; 2.07] | 0.58 [0.55; 0.72] | 0.13* [0.00; 1.46] | 0.02 [-0.03; 0.12] |
| fr (min<sup>-1</sup>) | 17.9 [15.2; 19.3] | 10.8 [4.0; 15.4] | 15.3 [9.4; 18.7] | -7.5*** [-10.9; -4.6] | -1.5*** [-4.8; -0.6] |
| Ti (s) | 1.4 [1.3; 1.6] | 1.7 [1.2; 2.1] | 1.6 [1.3; 1.7] | 0.3 [0.0; 0.6] | 0.1 [0.0; 0.2] |
| TE (s) | 1.7 [1.6; 2.2] | 2.3 [1.6; 2.9] | 2.1 [1.7; 3.0] | 0.3* [-0.2; 0.9] | 0.1 [0.0; 0.8] |
| Tp<sub>2</sub> (s) | 0.1 [0.0; 0.2] | 0.3 [0.1; 9.5] | 0.1 [0.1; 0.7] | 0.2*** [0.0; 9.3] | 0.1*** [0.0; 0.5] |
| Ptc<sub>CO<sub>2</sub></sub> (mmHg) | 41.7 [40.5; 45.8] | 42.0 [38.5; 44.2] | 41.4 [39.0; 44.1] | -1.4* [-3.3; 0.4] | -0.8* [-2.5; -0.2] |
| Pb<sub>CO<sub>2</sub></sub> (mmHg) | 42.0 [40.0; 44.0] | 41.8 [40.0; 44.3] | 44.0 [43.0; 44.8] | -0.3 [-1.0; 2.0] | 1.0 [0.0; 4.0] |
|  | Series: Immersion-RECOVERY SITTING |  |  |  |  |
| PET<sub>CO<sub>2</sub></sub> (mmHg, BTPS) | 30.2 [27.2; 33.2] | 32.2 [31.2; 34.1] | 29.1 [28.4; 31.0] | 2.3* [0.1; 4.3] | -0.6 [-2.0; 0.7] |
| PET<sub>O<sub>2</sub></sub> (mmHg, BTPS) | 111.4 [107.9; 114.9] | 110.2 [107.2; 113.1] | 111.5 [109.0; 113.5] | -1.7 [-3.5; 1.4] | 0.1 [-1.8; 1.7] |
| R | 0.817 [0.781; 0.891] | 0.903 [0.876; 0.970] | 0.820 [0.786; 0.846] | 0.089*** [0.068; 0.101] | -0.012 [-0.097; 0.039] |
| $\dot{V}_{CO_2}$ (l/min, STPD) | 0.24 [0.22; 0.26] | 0.23 [0.22; 0.26] | 0.24 [0.19; 0.28] | 0.00 [-0.03; 0.03] | 0.00 [-0.04; 0.03] |
| $\dot{V}_{O_2}$ (l/min, STPD) | 0.29 [0.24; 0.31] | 0.25 [0.23; 0.28] | 0.27 [0.25; 0.32] | -0.02 [-0.06; 0.01] | 0.00 [-0.01; 0.03] |
| MV (l/min, BTPS) | 10.0 [8.4; 12.0] | 9.0 [8.0; 9.7] | 9.9 [8.2; 11.5] | -1.4* [-2.9; -0.2] | 0.0 [-0.8; 0.7] |
| VT (l, BTPS) | 0.59 [0.50; 0.73] | 0.77 [0.60; 2.00] | 0.54 [0.49; 0.70] | 0.29** [0.07; 1.33] | 0.02 [-0.04; 0.04] |
| fr (min<sup>-1</sup>) | 18.5 [12.9; 21.7] | 12.0 [4.6; 15.7] | 16.0 [13.6; 20.2] | -6.1*** [-8.5; -4.5] | -0.9 [-2.5; 0.9] |
| Ti (s) | 1.3 [1.1; 1.4] | 1.3 [1.1; 1.5] | 1.3 [1.2; 1.4] | 0.0 [-0.2; 0.3] | 0.0 [-0.1; 0.0] |
| TE (s) | 1.8 [1.5; 2.4] | 1.7 [1.4; 2.9] | 1.9 [1.6; 2.2] | -0.2 [-0.7; 0.3] | 0.0 [-0.2; 0.3] |
| Tp<sub>2</sub> (s) | 0.2 [0.1; 0.5] | 0.7 [0.1; 7.5] | 0.4 [0.1; 1.0] | 0.7** [0.0; 7.7] | 0.1 [0.0; 0.7] |
| Ptc<sub>CO<sub>2</sub></sub> (mmHg) | 38.6 [37.5; 42.7] | 37.6 [34.8; 41.9] | 37.7 [35.3; 42.3] | -1.3** [-3.8; -0.8] | -0.7* [-2.4; 0.2] |
| Pb<sub>CO<sub>2</sub></sub> (mmHg) | 39.0 [39.0; 41.3] | 40.8 [38.5; 43.3] | 43.0 [41.5; 44.5] | 0.8 [-2.3; 2.0] | 4.0*** [2.5; 4.0] |
|  | Series: Immersion-CONTROL |  |  |  |  |
| PET<sub>CO<sub>2</sub></sub> (mmHg, BTPS) | 30.7 [29.6; 33.9] | 30.2 [29.3; 33.2] | 30.3 [28.8; 32.4] | -0.5* [-0.9; -0.3] | -1.1*** [-1.5; -0.4] |
| PET<sub>O<sub>2</sub></sub> (mmHg, BTPS) | 109.9 [107.8; 112.9] | 110.9 [108.9; 112.6] | 112.0 [108.6; 112.3] | 0.4 [-0.2; 1.4] | 0.5 [0.1; 2.2] |
| R | 0.846 [0.789; 0.927] | 0.839 [0.817; 0.903] | 0.838 [0.803; 0.901] | -0.005 [-0.024; 0.013] | -0.027 [-0.047; 0.040] |
| $\dot{V}_{CO_2}$ (l/min, STPD) | 0.23 [0.22; 0.28] | 0.22 [0.20; 0.24] | 0.23 [0.19; 0.24] | -0.03* [-0.03; 0.00] | -0.04 [-0.04; 0.00] |
| $\dot{V}_{O_2}$ (l/min, STPD) | 0.28 [0.25; 0.31] | 0.26 [0.24; 0.29] | 0.26 [0.23; 0.28] | -0.01* [-0.03; 0.00] | -0.02* [-0.04; -0.01] |
| MV (l/min, BTPS) | 9.5 [8.8; 11.2] | 9.0 [8.7; 11.0] | 9.9 [8.7; 10.6] | -0.3 [-0.8; 0.1] | -0.8 [-1.1; 0.3] |
| VT (l, BTPS) | 0.63 [0.58; 0.72] | 0.61 [0.54; 0.70] | 0.63 [0.53; 0.70] | 0.00 [-0.06; 0.01] | -0.02 [-0.07; 0.03] |
| fr (min<sup>-1</sup>) | 15.9 [14.6; 16.8] | 16.0 [14.6; 17.1] | 15.7 [14.8; 16.8] | 0.1 [-0.5; 0.4] | -0.5 [-0.9; 0.6] |
| Ti (s) | 1.5 [1.4; 1.6] | 1.5 [1.5; 1.7] | 1.6 [1.5; 1.6] | 0.0 [0.0; 0.1] | 0.0 [-0.1; 0.1] |
| TE (s) | 2.0 [1.8; 2.3] | 2.0 [1.9; 2.4] | 2.1 [2.0; 2.4] | 0.0 [-0.2; 0.1] | 0.1 [0.0; 0.2] |
| Tp<sub>2</sub> (s) | 0.1 [0.0; 0.1] | 0.1 [0.0; 0.1] | 0.1 [0.0; 0.1] | 0.0 [0.0; 0.0] | 0.0 [0.0; 0.0] |
| Ptc<sub>CO<sub>2</sub></sub> (mmHg) | 43.4 [40.7; 45.3] | 43.0 [42.2; 45.9] | 43.1 [41.1; 44.8] | 0.9 [-0.4; 1.6] | 0.2 [-1.5; 1.0] |
| Pb<sub>CO<sub>2</sub></sub> (mmHg) | 41.0 [38.8; 43.3] | 44.0 [42.4; 47.1] | 44.0 [43.0; 46.5] | 3.0*** [1.8; 4.1] | 3.0*** [2.0; 4.3] |

The results are presented as M [Q1; Q3], M is the median (calculated over the group of volunteers), Q1 and Q3 are first and third quartiles. Parameters designations are given in the text; \* - p<0.05, \*\* - p<0.01, \*\*\* - p<0.005 are given for pairwise comparison of “NPBin” stage or “after NPBin” stage with stage “before NPBin” of the current series (Wilcoxon's signed rank test).

Almost all parameters significantly (p<0.05) responded to NPBin in most series (excluding controls), except 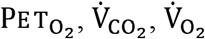, T_I_.

No pauses from inspiration to subsequent expiration were observed during free breathing and under NPBin (Tp_1_ = 0). Pauses between expiration and subsequent inspiration were absent during free breathing in the majority of volunteers. When switching to NPBin, pauses from expiration to subsequent inspiration appeared or increased significantly.

Under NPBin, V_T_ increased (by an average of 0.3 l) and f_R_ decreased (by an average of 7 min^-1^, up to 2-3 cycles per minute in some volunteers). The decrease in f_R_ was caused primarily by the increase in Tp_2_. Under NPBin, T_I_ and T_E_ slightly increased, which, apparently, is due to an increase in V_T_, for expiration the effect was more pronounced.

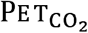 and 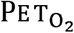changed in opposite directions under NPBin ( 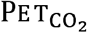increased by an average of 3 mmHg, 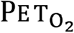 decreased by an average of 2 mmHg), MV slightly decreased (by an average of 1.8 l/min), and R increased by an average of 0.09.

Pairwise comparison of the same stages of HDBR-NPBin and HDBR-CONTROL series using Wilcoxon signed rank test did not reveal significant differences between “before NPBin” and “after NPBin” stages (p>0.05). For the “NPBin” stage, significant differences were observed for all parameters, except for 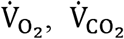, and MV. A similar pattern was observed when comparing the Immersion-CONTROL series with the Immersion-BDC SUPINE, Immersion-RECOVERY SUPINE, and Immersion-DI series. Therefore, NPBin effects on respiration and gas exchange did not persist after the end of NPBin.

When comparing changes in parameters relative to their initial value at the same stages of different series of DI experiment, no significant differences in the response to NPBin were found. Change in body position or DI did not affect the response to NPBin.

### 3.2 Transcutaneous data

Among the parameters obtained with the help of transcutaneous monitor, in all series significant changes (p<0.05) were observed only for 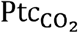 . Transcutaneous CO_2_ tension slightly decreased under NPBin (by 1.0-3.3 mmHg) and often remained reduced after returning to free breathing mode.

When comparing changes in parameters relative to their initial value at the same stages of DI experiment series with actual NPBin, we found no effects of body position or DI on the NPBin-induced decrease in 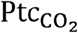 (p>0.05).

### 3.3 Capillary blood data

CO_2_ tension in arterialized capillary blood 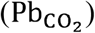 slowly increased in any of the series, the increase between the measurements performed at the beginning and at the end of a series was approximately 2-3 mmHg. In most series, including control ones, the increase was statistically significant (p<0.05). The reason for these changes in 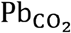 remains unclear, and they mask the effect of NPBin. However, the magnitude of the increase in 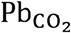 between the stages “NPBin” and “before NPBin” in control series is higher. In addition, in some cases, a decrease in 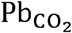 was observed under NPBin. Individual 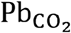 changes (“NPBin” stage minus “before NPBin” stage) obtained in all series were compared with changes observed in control series. There were significant (p<0.05) differences between Immersion-CONTROL and Immersion-BDC SUPINE series and between Immersion-CONTROL and Immersion-RECOVERY SUPINE series. Comparisons of Immersion-CONTROL series with Immersion-DI series and HDBR-NPBin with HDBR-CONTROL series did not reveal any significant differences. In a similar comparison of stages “after NPBin”, no significant differences were found for the mentioned above pairs of series. Apparently, NPBin leads to a slight decrease in 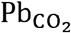 (within a few mmHg), and the decrease overlaps with the opposite drift of 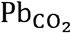 which is not associated with NPBin.

We found no significant changes in 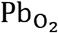 during any of the series. Dry immersion and changes in body position did not affect 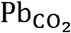 and 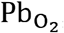, as well as 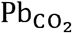 changes, during any of the series.

## 4. Discussion

Breathing pattern changes profoundly under NPBin. Probably, the respiratory system minimizes the additional load on respiratory muscles by decreasing f_R_ without increasing duration of inspiration.

It is worth noting the difference between the maximum rarefaction during inspiration under NPBin and the averaged over the entire “NPBin” stage value of Pm. If we assume that the average Pm value is the measure of NPBin impact on the cardiorespiratory system (on average, about -5 cmH_2_O), it turns out to be 4-5 times less than expected if we rely only on the settings of the valve that implements NPBin (-20 cmH_2_O). That happens due to increase in the duration of respiratory cycle under NPBin with almost no increase in the duration of inspiration. So increasing the portion of inspiration in the respiratory cycle may be considered as a way to increase the response of cardiorespiratory system to NPBin, or it is better to use NPB instead of NPBin.

The observed NPBin-induced changes in 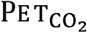 may be caused by the increase in V_T_, so the composition of end-expired portion of gas approximated the composition of alveolar gas, or they may be caused by changes in the efficiency of gas exchange in the lungs under NPBin. The NPBin-induced increase in R may reflect an increase in the work of respiratory muscles or an increase in the efficiency of gas exchange. But the decrease in 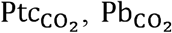 data, and the increase in R indicate rather a change in the efficiency of gas exchange in the lungs. The persistence of 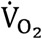 and 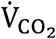 together with the decrease in MV under NPBin also indicates an increase in the efficiency.

Increased gas exchange efficiency leads to some washing out of CO_2_ from the body, despite the decrease in f_R_ (in some volunteers up to two cycles per minute!). The increase in R and 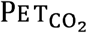 under NPBin is consistent with the results of transcutaneous gas tension measurements and capillary blood data, which demonstrates a slight decrease in 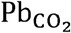and 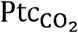 induced by NPBin. In addition, a slight increase in SpO_2_ has been observed under NPBin (Semenov et al., 2014), which is also indirectly indicates an increase in the efficiency of gas exchange in the lungs.

The increase in the efficiency is likely due to a decrease in the unevenness of ventilation and perfusion of the lungs, as well as due to the intensification of pulmonary circulation under NPBin. The increase can be associated both with the effect of decreased intrathoracic pressure and with changes in breathing pattern (an increase in V_T_). A special study is required to separate the contributions of the decrease in intrathoracic pressure under NPBin and the change in the pattern itself (without additional rarefaction).

The possible increase in gas exchange efficiency under NPBin deserves particular attention, as it may be desirable in some situations, such as in the treatment of respiratory diseases, or may lead to excessive CO_2_ washout affecting circulation.

## 5. Conclusions

NPBin-induced changes in respiration and gas exchange parameters were studied in detail for the first time under conditions of dry immersion and head-down bed rest.

Parameters of respiration and gas exchange responded to NPBin in a standard way under any of the considered experimental conditions (body positions, DI, HDBR): tidal volume increased, respiratory rate decreased, respiratory minute volume slightly reduced or remained unchanged, while end-tidal CO_2_ concentration and respiratory exchange ratio increased. Changes in respiratory rate were associated almost entirely with an increase (or appearance) of the pause between expiration and subsequent inspiration; other phases of respiratory cycle remained virtually unchanged when switching to NPBin. Changes in gas exchange parameters were accompanied by a slight decrease in CO_2_ tension in tissues and in arterialized capillary blood, other parameters did not respond significantly to NPBin.

NPBin effects on respiratory and gas exchange parameters did not persist after the end of NPBin (parameters returned to their original values within 1-5 minutes). We found no significant differences in NPBin-induced changes of the parameters between various experimental conditions (HDBR, DI, body positions).

The specified set of changes in respiratory and gas exchange parameters caused by NPBin indicates a washout of CO_2_ from tissues of the body under NPBin, despite the decrease in lungs minute ventilation and the increase in the work of breathing. This allows us to suppose that NPBin increases the efficiency of gas exchange in the lungs.

It is also worth noting that due to significant individual variations in the magnitude of respiratory rate decrease under NPBin and due to almost unchanged duration of inspiration, the average airway pressure under NPBin is also subject to significant individual scatter, even if there are no variations in peak rarefaction during inspiration. Besides, due to the decrease in respiratory rate under NPBin, average rarefaction in airways is significantly reduced relative to the expected value or to the additional rarefaction in airways during inspiration under NPBin. This phenomenon apparently weakens the effect of NPBin on the respiratory system and circulation, especially when considering slow processes like tissue gas diffusion or changes in tissue hydration. Possible diminution of NPBin effects caused by the decrease in respiratory rate should be taken into account when studying physiological responses to NPBin.

## Acknowledgments

The authors express their sincere gratitude for the friendly support and for the opportunity to take part in the experiments to the staff of the Department of Sensorimotor Physiology and Countermeasure of the Institute of Biomedical Problems of the Russian Academy of Sciences, who organized the dry immersion experiment, and to the staff of the Laboratory of Nutrition, Gastroenterology and Hygienic Control of Physical Environmental Factors of the Institute of Biomedical Problems of the Russian Academy of Sciences, who organized the bed rest experiment, and personally thank Elena Tomilovskaya, Ph.D., and Dr. Boris Afonin, Ph.D.

This work was supported by the Russian Academy of Sciences, Program “Integration of Control in ensuring the functions of the body” (grant IV.7.1.) and by the Program of basic research of the State Scientific Center of the Russian Federation – Institute of Biomedical Problems of the Russian Academy of Sciences (grant FMFR-2024-0038).

The research was carried out at the unique scientific installation “Medical and technical complex for the development of innovative space biomedicine technologies for the purpose of ensuring orbital and interplanetary flights, as well as for the development of practical healthcare”.

